# Evolution of the SARS-CoV-2 mutational spectrum

**DOI:** 10.1101/2022.11.19.517207

**Authors:** Jesse D Bloom, Annabel C Beichman, Richard A Neher, Kelley Harris

## Abstract

SARS-CoV-2 evolves rapidly in part because of its high mutation rate. Here we examine whether this mutational process itself has changed during viral evolution. To do this, we quantify the relative rates of different types of single nucleotide mutations at four-fold degenerate sites in the viral genome across millions of human SARS-CoV-2 sequences. We find clear shifts in the relative rates of several types of mutations during SARS-CoV-2 evolution. The most striking trend is a roughly two-fold decrease in the relative rate of G→T mutations in Omicron versus early clades, as was recently noted by Ruis et al (2022). There is also a decrease in the relative rate of C→T mutations in Delta, and other subtle changes in the mutation spectrum along the phylogeny. We speculate that these changes in the mutation spectrum could arise from viral mutations that affect genome replication, packaging, and antagonization of host innate-immune factors—although environmental factors could also play a role. Interestingly, the mutation spectrum of Omicron is more similar than that of earlier SARS-CoV-2 clades to the spectrum that shaped the long-term evolution of sarbecoviruses. Overall, our work shows that the mutation process is itself a dynamic variable during SARS-CoV-2 evolution, and suggests that human SARS-CoV-2 may be trending towards a mutation spectrum more similar to that of other animal sarbecoviruses.

## Introduction

The evolution of SARS-CoV-2 is enabled in part by the high underlying rate at which mutations arise in the viral genome during replication. Coronaviruses are the only RNA viruses known to have a proofreading mechanism in their RNA-dependent RNA polymerase (Denison et al. 2011)—but despite that proofreading, coronaviruses still have mutation rates that dwarf those of cellular organisms by several orders of magnitude (Drake 1993; Peck and Lauring 2018).

Studies of cellular organisms ranging from bacteria to humans have shown that the mutational process itself can change during evolution (Hwang and Green 2004; Sung, Tucker, et al. 2012; Couce et al. 2013; Long et al. 2015). Many studies of changes in the mutational process during natural evolution focus on the mutation spectrum which represents the distribution of *relative* rather than absolute rates of different types of mutations. For instance, humans experienced a transient increase in the relative rate of C→T mutations in certain sequence contexts, which affected a 10,000-year-old population of Anatolian hunter gatherers and spread via gene flow to all living Europeans and South Asians (Harris 2015; Speidel et al. 2021). The mutation spectrum also diverged more gradually during human evolution in Africa and East Asia, resulting in profiles that are sufficiently distinctive to identify an individual’s continent of origin. Populations of great apes, mice, and yeast have similarly distinctive mutational processes (Lindsay, et al. 2019; Jiang et al. 2021; Goldberg and Harris 2022). It remains unclear how much these changes are due to evolution of the underlying genome-replication machinery versus changes in life history or exposure to environmental mutagens (Mathieson and Reich 2017; Macià et al. 2021; Ruis, Peacock, et al. 2022; Ruis, Weimann, et al. 2022) — although in a few cases changes in the mutation spectrum have been linked to heritable genetic change affecting the function or expression of proteins involved in genome replication or repair (Couce et al. 2013; Jiang et al. 2021; Robinson, et al. 2021; Kaplanis et al. 2022; Sasani et al. 2022). For viruses, the mutational process can also be affected by a virus’s ability to evade host innate-immune proteins that mutagenize viral nucleic acids (Sadler et al. 2010; De Maio et al. 2021; Ratcliff and Simmonds 2021; Ringlander et al. 2022).

For coronaviruses like SARS-CoV-2, genes encoding proteins involved in genome replication and innate-immune antagonism constitute a substantial fraction of the genome (Ziebuhr 2005; V’kovski et al. 2021)—providing an ample target for mutations that could potentially alter the mutation process itself. In artificial lab settings, researchers have isolated coronavirus variants with mutations in genome-replication proteins that have dramatically altered mutation rates (Eckerle et al. 2007; Eckerle et al. 2010). However, it is unclear how such mutator variants generally fare during natural evolution (Peck and Lauring 2018).

A recent preprint by Ruis, et al. on pathogenic bacterial mutagenesis identified several mutation types whose relative rates correlate with replication niche within the human body (Ruis, Weimann, et al. 2022). The authors found that bacterial replication within the lower respiratory tract correlated with an increased load of G→T mutations, which prompted them to hypothesize that the Omicron lineage of SARS-CoV-2 would have a reduced G→T rate relative to earlier SARS-CoV-2 lineages that may replicate more in the lungs (Ruis, Peacock, et al. 2022). Consistent with this hypothesis, they found a reduced relative number G→T mutations across all sites for Omicron clades of SARS-CoV-2. Since their study pooled all mutations (nonsynonymous and synonymous), it is not clear the extent to which the signal could be affected by protein-level selection as well as the underlying rate of mutation. It is also unclear whether the difference in G→T fraction between Omicron and other SARS-CoV-2 viruses is the dominant feature of the SARS-CoV-2 mutational landscape or just one component of the sort of continuous variation that has been observed in cellular organisms.

Here we systematically analyze changes in the relative rates of all single nucleotide mutation types among different clades of human SARS-CoV-2. To disentangle underlying mutation rates from the subsequent action of natural selection, we restrict our analysis to only four-fold degenerate sites where all mutations are expected to be neutral with respect to protein function. We also use rigorous quality-control to ensure our estimates are not biased by technical artifacts related to sequencing or base-calling errors. Using this approach, we confirm that Omicron has a roughly two-fold decrease in the relative rate of G→T mutations relative to early clades. We also find additional shifts in the mutation spectrum, including a decrease in C→T mutations in Delta and a broader correlation between mutation spectrum divergence and genetic divergence across the SARS-CoV-2 phylogeny. While our analysis does not determine the cause of the evolutionary shifts in SARS-CoV-2’s mutational spectrum, the pervasive and phylogenetically correlated nature of the shifts suggests viral mutations affecting genome replication, packaging, and innate-immune antagonism could play a role.

## Results

### Different clades of human SARS-CoV-2 have different mutation spectra

We focused our analysis on the roughly 6-million publicly available SARS-CoV-2 sequences in the pre-built UShER phylogenetic tree (Turakhia et al. 2021). Each of these sequences represents the consensus sequence of a virus that infected a human individual. We counted the occurrence of each mutation on the branches of the phylogenetic tree (Turakhia et al. 2021): these counts represent the number of *occurrences* of mutations, not how often the mutations are found in the final sequence alignment (in other words, a mutation that occurred once but then is shared in several sequences by common descent is only counted once). We tallied counts separately for each Nextstrain clade (Aksamentov et al. 2021), and used a variety of quality-control steps to remove sequences and sites likely to be affected by spurious mutations from sequencing or base-calling artifacts (see Methods).

Prior analyses of SARS-CoV-2 mutation rates have generally focused on all nucleotide mutations (Neher 2022; Ruis, Peacock, et al. 2022). However, many sites in the viral genome are under strong functional selection, and so the mutational patterns at those sites will represent the combined action of mutation and selection. We therefore focused our analysis only on four-fold degenerate sites (sites at the third position in codons where all three possible nucleotide mutations are synonymous), under the assumption that mutations at such sites will tend to be nearly neutral. The SARS-CoV-2 genome has ~4,240 such sites (with the exact number differing slightly among viral clades), and we restricted our analysis to only clades with at least 5,000 mutations at such sites (Table 1).

**Table 1.** Number of four-fold degenerate synonymous sites and total mutations at those sites for the clades analyzed here. Note we only analyzed clades with at least 5,000 mutations at four-fold degenerate sites.

| clade | four-fold degenerate sites | total mutations at these sites |
| --- | --- | --- |
| 20A | 4,247 | 17,202 |
| 20B | 4,247 | 14,121 |
| 20C | 4,246 | 9,344 |
| 20E | 4,246 | 10,454 |
| 20G | 4,243 | 14,019 |
| 20I | 4,243 | 60,858 |
| 21C | 4,245 | 6,308 |
| 21I | 4,241 | 24,117 |
| 21J | 4,239 | 282,051 |
| 21K | 4,241 | 113,721 |
| 21L | 4,235 | 83,475 |
| 22A | 4,236 | 11,413 |
| 22B | 4,234 | 64,765 |
| 22C | 4,233 | 18,958 |

There were clear differences in the mutation spectrum at four-fold degenerate sites across viral clades (Figure 1A and interactive plot at https://jbloomlab.github.io/SARS2-mut-spectrum/pca.html). The largest difference was between Omicron clades and all other clades, but Delta clades also showed a unique pattern. Importantly, these clade-to-clade differences were robust to analyzing sequences only from specific geographical locations, excluding the most heavily mutated sites, or analyzing each half of the viral genome separately (Figure S1 and https://jbloomlab.github.io/SARS2-mut-spectrum/).

**Figure 1.**
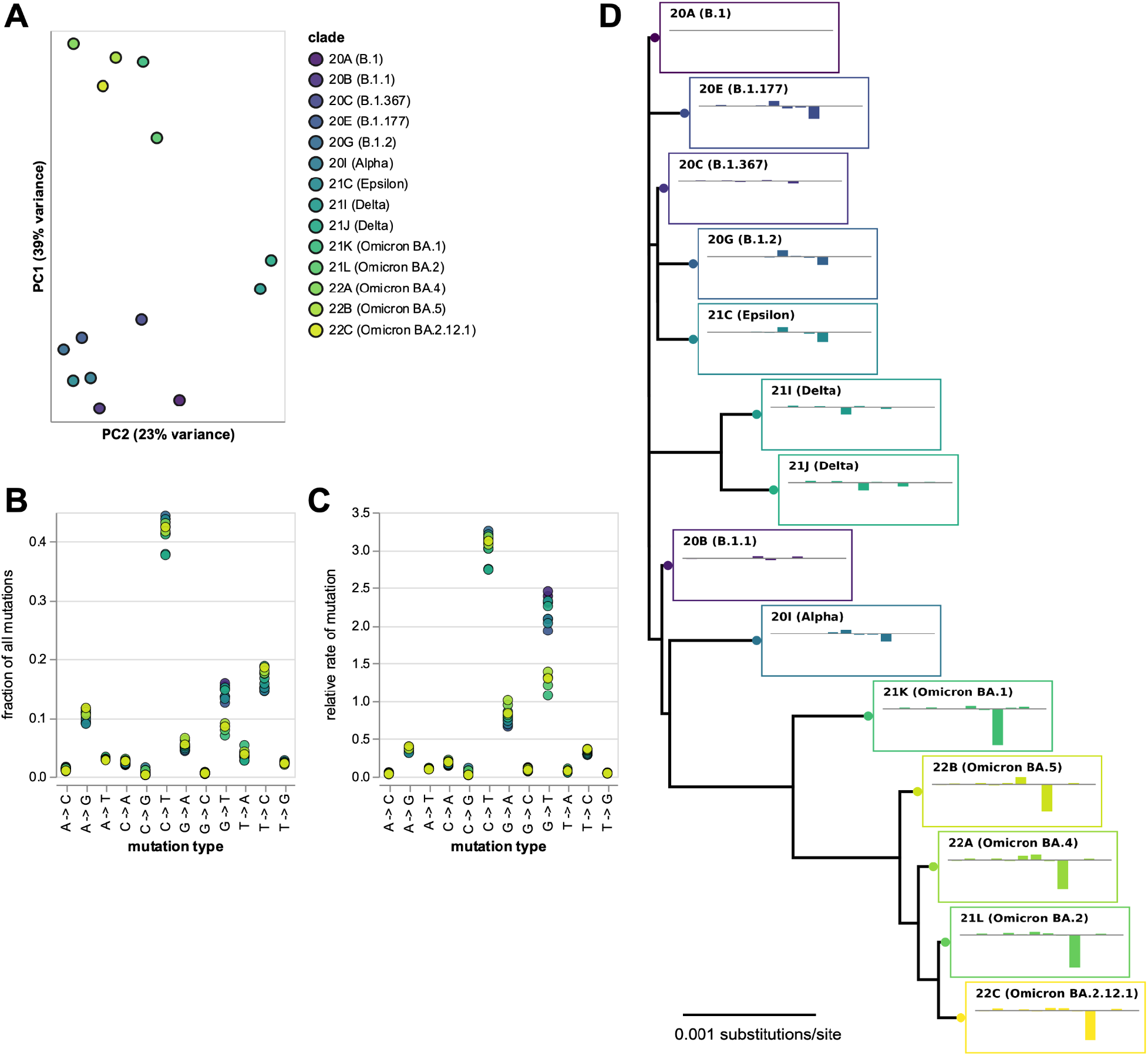
Mutation spectrum of SARS-CoV-2 clades at four-fold degenerate sites. **(A)** Principal component analysis (PCA) of mutation spectra of Nextstrain clades with sufficient sequences (each point is a clade). **(B)** Fraction of mutations at four-fold degenerate sites that are of each type for each clade. **(C)** Relative rates of each mutation type, calculated as the fraction of mutations of that type divided by the fraction of sites with the parental nucleotide. **(D)** Phylogenetic tree of clade founder sequences, with plots showing mutation rates for that clade (ordered as in panel C) minus rates for clade 20A. Interactive versions of panels A-C at https://jbloomlab.github.io/SARS2-mut-spectrum/ enable easier identification of individual clades. Figure S1 shows the PCA is robust to subsetting on sequences from different geographic locations, excluding top mutations, and partitioning the genome. The numerical values in panels (B) and (C) are at https://github.com/jbloomlab/SARS2-mut-spectrum/blob/main/results/synonymous_mut_rates/rates_by_clade.csv

The biggest difference between Omicron and other clades was a roughly two-fold decrease in the rate of G→T mutations (Figure 1B,C and interactive plot at https://jbloomlab.github.io/SARS2-mut-spectrum/rates-by-clade.html), consistent with a recent study (Ruis, Peacock, et al. 2022) that analyzed all sites (synonymous and nonsynonymous). There was also a clear decrease in the rate of C→T mutations in Delta (Figure 1B,C). Some other types of mutations with lower rates also showed substantial differences among clades (this is seen most easily by clicking on specific mutation types in the interactive plot at https://jbloomlab.github.io/SARS2-mut-spectrum/rates-by-clade.html). Note also that we confirm prior findings that the two types of mutations with the highest rates are C→T transitions and G→T transversions (De Maio et al. 2021).

### The mutation spectrum has phylogenetic signal beyond G→T mutations

The clade-to-clade differences in relative mutation rates has a visually obvious phylogenetic pattern (Figure 1D). To statistically confirm the visual impression of phylogenetic patterns in the mutation rates, we used Mantel tests (Mantel 1967; Harmon and Glor 2010; Hardy and Pavoine 2012; Legendre and Legendre 2012) to compare the phylogenetic distances between clades with the differences in their relative mutation rates (Figure 2). These tests showed that the relative mutation rates were indeed significantly correlated with the phylogenetic distances between clades. The correlations remained significant even if we excluded G→T mutations, or analyzed only Omicron or non-Omicron clades (Figure 2). These results show that evolution of the mutation spectrum goes beyond simply a change in the relative rate of G→T mutations in Omicron.

**Figure 2.**
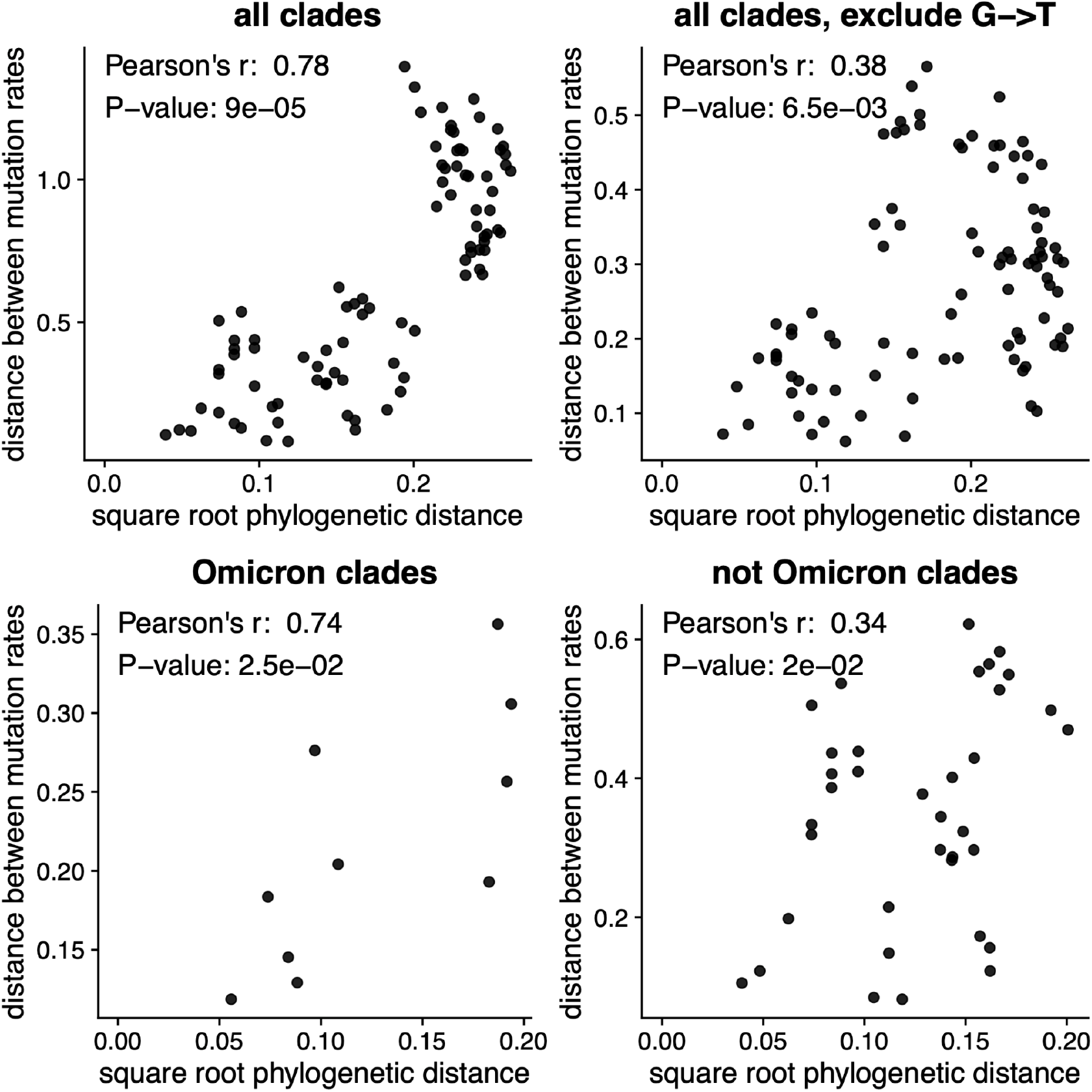
The changes in relative mutation rates correlate with the phylogenetic relationships among clades. The plots show the correlation between the square root of the phylogenetic distance separating each pair of clades and the Euclidean distance between the relative mutation rates for those clades. Assuming that mutation rates evolve neutrally according to a Brownian motion model, mutation rate distances should scale linearly with the square root of phylogenetic distance. The *p*-values are computed using the Mantel test with 100,000 permutations. The plots show that the mutation rates are correlated with phylogenetic distance even if we exclude G→T mutations, or do the analysis only among Omicron or non-Omicron clades.

The G→T mutation fraction change observed in Omicron (from an ancestral fraction of about 15% to a derived fraction of about 8%, see Figure 1B) could be the result of a 2-fold decrease in the absolute rate of G→T mutations in this lineage if the rates of all other mutations stayed approximately constant. Such a rate change would imply that the overall Omicron mutation rate is about 7% lower than the mutation rate of non-Omicron lineages. More complicated scenarios are also possible, such as an increase in the rates of all non-G→T mutation types in Omicron or compensatory increases and decreases of different mutation types that leave the overall rate unchanged. The existence of phylogenetic signal in the mutation spectrum of non-G→T mutations suggests that the rates of multiple mutation types likely changed over time, but none of these shifts necessarily imply a detectable change in the overall SARS-CoV-2 mutation rate.

### SARS-CoV-2’s mutation spectrum is becoming more similar to the mutation spectra of other sarbecoviruses

In the absence of natural selection, the nucleotide composition of a gene sequence should eventually reach a stable “equilibrium” nucleotide frequency distribution that is determined by its mutation spectrum (Felsenstein 2003). If we assume that the nucleotides at four-fold degenerate sites are not under selection, then the actual observed frequencies of nucleotides at these sites should be similar to the equilibrium frequencies predicted by the mutation spectrum if the virus has been evolving with the same mutation spectrum for a sufficiently long period of time.

We calculated the predicted equilibrium frequencies of nucleotides from the mutation spectra of the various human SARS-CoV-2 clades (Figure 3A). Because the mutation spectra differ somewhat among clades, the predicted equilibrium nucleotide frequencies also differ among clades: for instance, Omicron’s mutation spectrum implies a somewhat lower equilibrium frequency of T nucleotides than earlier clades, in part because Omicron has a lower rate of G→T mutations.

**Figure 3.**
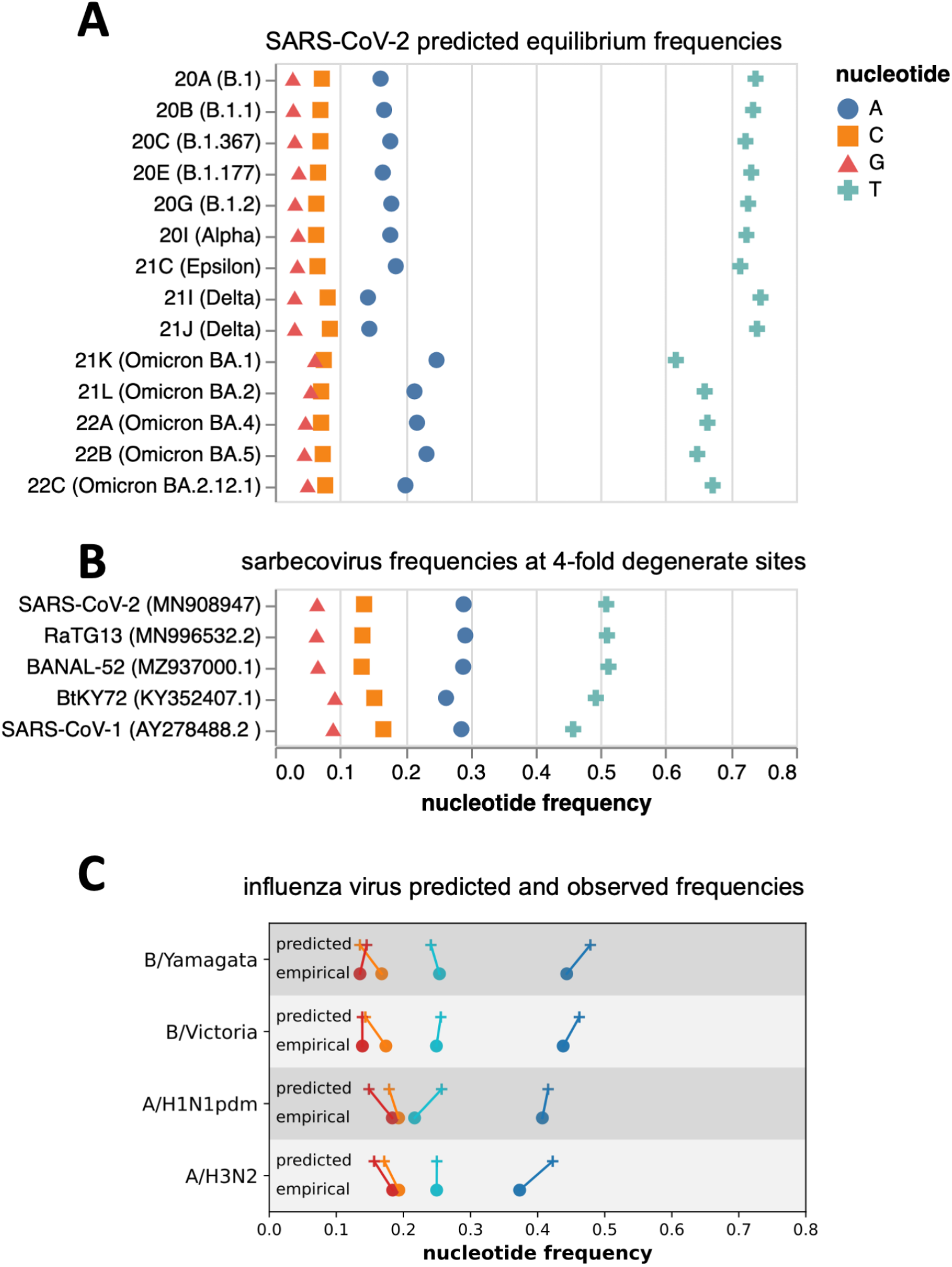
Predicted equilibrium frequencies from mutation rates versus actual nucleotide frequencies at four-fold degenerate sites in sarbecoviruses. (A) Predicted equilibrium frequencies of nucleotides at four-fold degenerate sites as calculated from the mutation rates for all of the SARS-CoV-2 clades analyzed here. (B) Actual frequencies of nucleotides at four-fold degenerate sites in various sarbecoviruses. (C) Predicted equilibrium frequencies (from mutation rates) and actual empirically observed nucleotide frequencies at four-fold degenerate sites for human influenza virus types and subtypes. The color code in panel C is the same as for panels A and B.

We compared these predicted equilibrium frequencies from the SARS-CoV-2 clades’ mutation spectra to the actual frequencies of nucleotides observed at four-fold degenerate sites in various sarbecoviruses (the subgenus of coronaviruses to which SARS-CoV-2 belongs). As can be seen from Figure 3B, the nucleotide frequencies at four-fold degenerate sites are similar among SARS-CoV-2, two close relatives (e.g., RaTG13 and BANAL-52), and more diverged sarbecoviruses such as SARS-CoV-1 and BtKY72, suggesting that the long-term evolution of all these viruses has been shaped by a similar mutation spectrum.

However, the equilibrium nucleotide frequencies predicted by the mutation spectrum of human SARS-CoV-2 are quite different from the actual frequencies observed in SARS-CoV-2 and other sarbecoviruses (Figure 3A,B). Some of this difference could be due to natural selection even at four-fold degenerate sites, or flanking context dependence in mutation rates not incorporated into our analysis. However, when we performed a similar analysis for human influenza virus, we found the empirically observed nucleotide frequencies at four-fold degenerate sites to be largely concordant with the equilibrium frequencies predicted from the influenza virus mutation spectrum (Figure 3C)—indicating that at least for some human RNA respiratory viruses, the observed nucleotide frequencies are compatible with the viral mutation spectrum.

The actual observed nucleotide frequencies of both SARS-CoV-2 and other sarbecoviruses are more similar to the equilibrium nucleotide frequencies implied by the mutation spectra of the Omicron clades are more similar than the frequencies implied by the spectra of earlier human SARS-CoV-2 clades (note how the Omicron clades in Figure 3A look more similar to Figure 3B). The long-term evolution of all these sarbecoviruses occurred in bats, and it is possible that some aspect of replication in humans altered the mutation spectrum of SARS-CoV-2, and is now shifting in Omicron back to a spectrum more similar to that of bat coronaviruses.

### Putative associations of protein-coding mutations with changes in the mutation spectrum

Mutation spectrum changes could potentially be caused by clade-specific amino-acid mutations in viral proteins involved in genome replication, packaging, or antagonization of host-cell innate-immune proteins that mutagenize foreign nucleic acids (De Maio et al. 2021; Ratcliff and Simmonds 2021; Ringlander et al. 2022). To explore the plausibility of this hypothesis, we tabulated the non-spike amino-acid mutations in each clade relative to the early 20A (B.1) clade (Table 2) and identified several viral amino-acid mutations that we speculate could potentially affect the mutation spectrum.

**Table 2.** Non-spike amino-acid mutations in the founder sequence for each clade relative to the early 20A (B.1) clade.

| clade | non-spike amino-acid mutations relative to clade 20A |
| --- | --- |
| 20B (B.1.1) | N: R203K, G204R |
| 20C (B.1.367) | ORF3a: Q57H; nsp2: T85I |
| 20E (B.1.177) | N: A220V; ORF10: V30L |
| 20G (B.1.2) | N: P67S, P199L; ORF3a: Q57H, G172V; ORF8: S24L; nsp14: N129D; nsp16: R216C; nsp2: T85I; nsp5: L89F |
| 20I (Alpha) | N: D3L, R203K, G204R, S235F; ORF8: Q27*, R52I, Y73C; nsp3: T183I, A890D, I1412T |
| 21C (Epsilon) | N: T205I; ORF3a: Q57H; nsp13: D260Y; nsp2: T85I |
| 21I (Delta) | M: I82T; N: D63G, R203M, D377Y; ORF3a: S26L; ORF7a: V82A, T120I; nsp12: G671S; nsp13: P77L; nsp3: P822L; nsp4: A446V; nsp6: V149A, T181I |
| 21J (Delta) | M: I82T; N: D63G, R203M, G215C, D377Y; ORF3a: S26L; ORF7a: V82A, T120I; ORF7b: T40I; nsp12: G671S; nsp13: P77L; nsp14: A394V; nsp3: A488S, P1228L, P1469S; nsp4: V167L, T492I; nsp6: T77A |
| 21K (Omicron BA.1) | E: T9I; M: D3G, Q19E, A63T; N: P13L, R203K, G204R; nsp14: I42V; nsp3: K38R, A1892T; nsp4: T492I; nsp5: P132H; nsp6: I189V |
| 21L (Omicron BA.2) | E: T9I; M: Q19E, A63T; N: P13L, R203K, G204R, S413R; ORF3a: T223I; ORF6: D61L; nsp1: S135R; nsp13: R392C; nsp14: I42V; nsp15: T112I; nsp3: T24I, G489S; nsp4: L264F, T327I, L438F, T492I; nsp5: P132H |
| 22A (Omicron BA.4) | E: T9I; M: Q19E, A63T; N: P13L, P151S, R203K, G204R, S413R; ORF3a: T223I; ORF6: D61L; ORF7b: L11F; nsp1: S135R; nsp13: R392C; nsp14: I42V; nsp15: T112I; nsp3: T24I, G489S; nsp4: L264F, T327I, T492I; nsp5: P132H |
| 22B (Omicron BA.5) | E: T9I; M: D3N, Q19E, A63T; N: P13L, R203K, G204R, S413R; ORF3a: T223I; nsp1: S135R; nsp13: R392C; nsp14: I42V; nsp15: T112I; nsp3: T24I, G489S; nsp4: L264F, T327I, T492I; nsp5: P132H |
| 22C (Omicron BA.2.12.1) | E: T9I; M: Q19E, A63T; N: P13L, R203K, G204R, S413R; ORF3a: T223I; ORF6: D61L; nsp1: S135R; nsp13: R392C; nsp14: I42V; nsp15: T112I; nsp3: T24I, G489S; nsp4: L264F, T327I, L438F, T492I; nsp5: P132H |

The Omicron clades all share mutation I42V in nsp14 (also known as ExoN), which provides proofreading activity during genome replication (Denison et al. 2011; Ogando et al. 2020). Rare polymerase proofreading defects have recently been shown to perturb the G→T mutation rate in human cells (Robinson, et al. 2021). The Omicron clades also share mutation P13L in the nucleoprotein, which is part of the genome replication complex and encapsidates viral RNA (Bessa et al. 2022), and P132H in nsp5, which proteolytically processes the polyprotein encoding the viral replicase (Roe et al. 2021) and helps antagonize innate immune responses (Liu et al. 2021). The Delta clades share mutation G671S in the viral polymerase nsp12 (Kirchdoerfer and Ward 2019), as well as several mutations in the nucleoprotein and a mutation in the ORF3a protein that may play a peripheral role in viral replication (Zhang et al. 2022). The Delta clades also share mutations in the ORF7a (V82A and T120I) and nsp13 (P77L) proteins involved in innate-immune antagonization (Cao et al. 2021; Fung et al. 2022)—which could be relevant as Delta has a decreased relative rate of the C→T mutations, which is the type of change induced by host-cell APOBEC innate-immune proteins (De Maio et al. 2021; Ratcliff and Simmonds 2021). Note also that non-coding mutations or indels (which are not listed in Table 1) could also affect the mutation spectrum if they alter expression of viral genes.

## Discussion

We have demonstrated that there are clear shifts in the mutation spectrum during the evolution of SARS-CoV-2. We corroborate the findings of Ruis, et al. that Omicron has a lower relative rate of G→T mutations (Ruis, Peacock, et al. 2022), but we also show that the changes in the mutation spectrum are not restricted to this one type of mutation. Instead, there are also significant phylogenetically correlated shifts in the spectrum among other mutation types, and among both Omicron and non-Omicron clades. In this sense, changes in the SARS-CoV-2 mutation spectrum appear to involve the type of pervasive evolutionary shifts that have been observed among many cellular organisms (Harris 2015; Lindsay, et al. 2019; Jiang et al. 2021; Goldberg and Harris 2022).

Our analysis cannot determine why the mutation spectrum differs among clades—although our approach of calculating the rates at only four-fold degenerate sites does rule out confounding effects of protein-level selection. Ruis et al proposed that the lower rate of G→T mutations in Omicron is due to a shift to viral replication in the upper rather than lower airways (Ruis, Peacock, et al. 2022). This is certainly possible, but we suggest that the differences may instead be driven by mutations elsewhere in the viral genome. For instance, Omicron and Delta have clade-specific mutations in proteins involved in genome replication, packaging, and innate-immune antagonism. The latter factor could be important since some mutations (such as the C→T mutations that are relatively rarer in Delta) can be induced by host-cell innate-immune proteins (De Maio et al. 2021; Ratcliff and Simmonds 2021). Ruis et al. accurately point out that Omicron does not have any amino-acid mutations in the active site of core genome replication proteins, but prior work has shown that the mutation rates of other viruses can be modulated by mutations distant from polymerase protein active sites (Vignuzzi et al. 2008; Pauly et al. 2017). Similar subtle modifications could be induced by mutations to the nucleoprotein (which is part of the replication complex and protects viral RNA), as well as proteins that modulate expression of host-cell innate-immune proteins. However, ultimate determination of the cause of the changes in the mutation spectrum will require experimental work beyond the scope of our study, and could also potentially be due to a wide range of factors including modifications in the location or speed of replication or transmission.

Our analysis examines the *relative* rather than *absolute* rates of different types of nucleotide mutations across SARS-CoV-2 clades. We take this approach because relative mutation rates can be internally calibrated, whereas precise estimation of absolute mutation rates from natural sequence data is harder. However, other recent work suggests that the overall absolute mutation rate is fairly similar among human SARS-CoV-2 clades (Neher 2022). But if the two-fold drop in the relative rate of G→T mutations in Omicron reflects a two-fold drop in absolute rate of that mutation type, that would only decrease the absolute rate across all mutations by ~7%, which would not be detectable at the resolution of current studies (Neher 2022). Note that much more dramatically elevated mutation rates have been observed in rare clusters of human (Hisner 2022) or white-tail deer SARS-CoV-2 (Pickering et al. 2022), but these clusters have not spread widely. Overall, these observations are consistent with the idea that mutation rates might drift moderately during natural evolution of successful SARS-CoV-2 variants (Sung, Ackerman, et al. 2012). However, so far there is no evidence for widespread transmission of SARS-CoV-2 variants with extreme changes in mutation rates like those sometimes observed in the lab (Eckerle et al. 2007; Eckerle et al. 2010; Pauly et al. 2017).

Interestingly, the actual nucleotide frequencies at four-fold degenerate sites in both SARS-CoV-2 and related sarbecoviruses differ from what would be predicted based on the mutation spectrum of any human SARS-CoV-2 clade. This difference is especially large for the mutation spectrum of early SARS-CoV-2 clades, with the mutation spectrum of Omicron clades being closer to that which shaped the long-term evolution of sarbecoviruses. We acknowledge that comparison of the mutation rates estimated in our study to nucleotide frequencies in natural sarbecoviruses could be somewhat confounded if there is weak selection on nucleotide identity even at four-fold synonymous sites. But we were able to confirm that four-fold degenerate nucleotide frequencies are close to their expected equilibrium for influenza A and B viruses, suggesting SARS-CoV-2 may be unusual in having a mutation spectrum that is highly discordant with the actual frequencies of nucleotides at putatively neutral sites. One possible explanation is that the mutation spectrum of sarbecoviruses could be relatively stable in the natural reservoir of bats, but has been altered in SARS-CoV-2 by some aspect of replication in humans—and is now undergoing relatively rapid evolutionary change.

The broader implications of shifts in the mutation spectrum of SARS-CoV-2 for its evolution are unclear. Changes in the mutation spectrum alter the rates at which different potentially adaptive amino-acid mutations arise. But SARS-CoV-2 evolution in humans exhibits high levels of convergence (Martin et al. 2021; Cao et al. 2022), with putatively beneficial amino-acid mutations often emerging many independent times in different viral variants. This convergence suggests that the virus’s evolution is not generally limited by the underlying rate at which new mutations appear. Therefore, the changes in the mutation spectrum we report are likely to at most modestly impact the overall process of adaptive evolution. However, our work does suggest that clade-specific estimates of the mutation rate are likely to improve the accuracy of efforts to estimate the fitness effects of viral mutations from their number of observed occurrences in natural sequences (Neher 2022), and could perhaps be useful for certain types of phylogenetic analyses. In addition, our work shows that the mutation process is clearly dynamic during SARS-CoV-2 evolution, so it will be interesting to see if larger changes in the mutation spectrum accrue as the virus continues to evolve.

## Methods

### Counting mutations at four-fold degenerate sites

We determined the mutation spectrum by counting the number of unique occurrences of each nucleotide mutation on the branches of a global phylogenetic tree of all publicly available SARS-CoV-2 sequences. Note we are counting how many times each mutation is inferred to have independently *occurred* among available consensus SARS-CoV-2 sequences from individual human infections, not its final count in the alignment of such sequences (this distinction is important because a single occurrence of a mutation may be observed in multiple sequences due to shared ancestry).

To count mutations, we used the pre-built clade-annotated UShER mutation-annotated tree (Turakhia et al. 2021) from November-7-2022 (http://hgdownload.soe.ucsc.edu/goldenPath/wuhCor1/UShER_SARS-CoV-2/2022/11/07/public-2022-11-07.all.masked.nextclade.pangolin.pb.gz). We used matUtils (Turakhia et al. 2021) to subset the mutation-annotated tree on samples from each Nextstrain clade, and then extract the mutations on each branch of the subsetted mutation-annotated trees. We next tallied the counts of each mutation on all branches for that clade, excluding mutations on any branches with >4 total mutations, >1 mutation that was a reversion to either the Wuhan-Hu-1 reference genome (Genbank NC_045512.2), or >1 mutation that was a reversion to the founder for that Nextstrain clade as defined by (Neher 2022) (see https://github.com/neherlab/SC2_variant_rates/blob/62c525dc4238385ec0755b40658f3007fdbfab1a/data/clade_gts.json). The rationale for these exclusions is that branches with abnormally large numbers of mutations are often indicative of low-quality sequences with lots of errors, and branches with abnormally large numbers of reversions to the reference or clade founder can be indicative of sequences generated by problematic bioinformation pipelines that call low-coverage regions to the reference.

For each clade, we then identified sites that are four-fold degenerate in the clade founder (see https://github.com/jbloomlab/SARS2-mut-spectrum/blob/main/results/clade_founder_nts/clade_founder_nts.csv). We also manually excluded sites that previous analyses (Turakhia et al. 2020) or our own analysis suggested might be prone to errors due to abnormally large numbers of mutations (the excluded sites are listed under *sites_to_exclude* in https://github.com/jbloomlab/SARS2-mut-spectrum/blob/main/config.yaml). Finally, we excluded any sites that differed between the clade founder and the Wuhan-Hu-1 reference (ie, had fixed mutations in the clade founder relative to Wuhan-Hu-1). This exclusion was designed to avoid any spurious mutations caused by bioinformatics pipelines that call low-coverage sites to reference. The counts for *all* mutations in each clade are in the file at https://github.com/jbloomlab/SARS2-mut-spectrum/blob/main/results/mutation_counts/aggregated.csv which contains columns indicating which sites are four-fold degenerate or specified for exclusion. Table 1 shows the number of four-fold degenerate sites for each clade and the total number of mutations at these sites. Note we only retained clades with at least 5000 mutation counts at non-excluded four-fold degenerate sites.

Finally, we tabulated the counts for each type of nucleotide mutation for each clade at the non-excluded four-fold degenerate sites, and determined the fraction of all mutations that were of that type (https://github.com/jbloomlab/SARS2-mut-spectrum/blob/main/results/synonymous_mut_rates/rates_by_clade.csv).

For the analyses in Figure S1, we repeated the above process but subsetted only sequences from the USA or England (as determined by whether the strain name contained that word), after excluding any site that was among the top 5 most mutated sites for any clade, or after partitioning the genome into halves.

### Principal component analysis

The principal component analyses (PCAs) were performed on the length 12 probability vectors giving the fraction of all mutations at the four-fold degenerate sites that were of each mutation type. The PCA was done using *scikit-learn* after first standardizing the vectors to have zero mean and unit variance. As described above, we repeated this analysis on subsets of the data to determine whether the results remained consistent when we restricted our analyses to only sequences from the USA and England, excluded any site that was among the top 5 most mutated sites for any clade, or partitioned the genome into halves.

### Calculation of relative mutation rates

The relative mutation rates plotted in Figure 1C were calculated simply by normalizing the fraction of all four-fold degenerate mutations that were of a given type by the fraction of all nucleotides at those sites in the clade founder that were of the parental nucleotide identity. For instance, the relative rate of A→T mutations was computed as the fraction of all mutations at non-excluded four-fold degenerate sites that changed an A to a T, divided by the fraction of all four-fold degenerate sites that had an A as their identity in the clade founder. Note that the frequencies of the different nucleotides at four-fold degenerate sites is virtually identical among the clade founder sequences (Figure S2).

### Phylogenetic tree

The phylogenetic tree in Figure 1D was inferred on the clade founder sequences using *iqtree* (Minh et al. 2020), and then rendered using *ete3* (Huerta-Cepas et al. 2016). The tips show the relative rates (as in Figure 1C) for each clade minus those rates for clade 20A, with the mutation types in the same order as in Figure 1C.

### Mantel test

The Mantel test (Mantel 1967; Harmon and Glor 2010; Hardy and Pavoine 2012; Legendre and Legendre 2012) was used to estimate the significance of the correlation between the Euclidean distance between clades’ mutation spectra and the square root of the phylogenetic distance between clade founder sequences (as estimated using *iqtree*), also known as phylogenetic signal (Figure 2). The square root of the phylogenetic distance is used because it is expected to scale linearly with Euclidean distance under a Brownian motion model (Hardy and Pavoine 2012). The Mantel test was implemented using the R package vegan (version 2.5-7), with method=“pearson” and 100,000 permutations (Oksanen et al. 2022). To determine whether the phylogenetic signal we observe is solely due to Omicron’s G>T fraction, the Mantel test was also carried out after excluding G>T mutations from the mutation spectrum, and re-normalizing it. To additionally determine whether the phylogenetic signal is due only to differences between Omicron and non-Omicron clades, we also carried out tests for phylogenetic signal on Omicron clades and non-Omicron clades separately.

### Equilibrium frequencies of SARS-CoV-2 nucleotides

The predicted equilibrium frequencies of nucleotides shown in Figure 3A were calculated as the real component of the principal eigenvector of a rate matrix constructed from the relative rates of each mutation type for that clade.

### Predicted and actual nucleotide frequencies at four-fold degenerate sites for influenza virus

The predicted and observed nucleotide frequencies for human influenza virus in Figure 3C were calculated from phylogenetic analyses of the different influenza virus lineages at nextstrain.org/groups/neherlab and influenza reference genomes used in the nextstrain seasonal influenza workflow. The empirical nucleotide frequencies were calculated by counting nucleotide states at four-fold synonymous sites in genomes of A/Beijing/32/1992 (A/H3N2), A/California/07/2009 (H1N1pdm), B/Hong Kong/02/1993 (B/Vic), and B/Singapore/11/1994 (B/Yam). The mutation spectrum was calculated by traversing the phylogenetic tree and counting mutations at positions that are four-fold synonymous in the reference sequence. From the spectrum and the empirical equilibrium frequencies, the predicted equilibrium frequencies were calculated as for SARS-CoV-2. The six largest segments PB2, PB1, PA, HA, NP, and NA were used for these analyses. These calculations are explicitly documented in the https://github.com/jbloomlab/SARS2-mut-spectrum/blob/main/scripts/compare_flu_spectra.py script A table listing the originating and submitting labs of influenza sequences used in this analysis is provided at https://github.com/jbloomlab/SARS2-mut-spectrum/blob/main/GISAID_acknowledgments/flu_acknowledgement.tsv

## Data and code availability

The computer code used for the analysis is available at https://github.com/jbloomlab/SARS2-mut-spectrum as a fully reproducible *Snakemake* pipeline (Mölder et al. 2021). Interactive versions of many of the plots rendered with *Altair* (VanderPlas et al. 2018) are at https://jbloomlab.github.io/SARS2-mut-spectrum/

## Acknowledgments

We thank Ryan Hisner and Adam Lauring for helpful comments. This research is based on sequence data by hundreds of laboratories around the world that have generously shared their data. We gratefully acknowledge their contributions. This work was supported in part by the NIH/NIAID grant R01AI141707 to JDB, NIH/NIA T32AG066574 to ACB, NIH/NIGMS grant R35GM133428 to KH, a Burroughs Wellcome Career Award at the Scientific Interface to KH, a Searle scholarship to KH, a Pew Scholarship to KH, and a Sloan Fellowship to KH. JDB is an Investigator of the Howard Hughes Medical Institute.

## Supplementary Material

**Figure S1.**
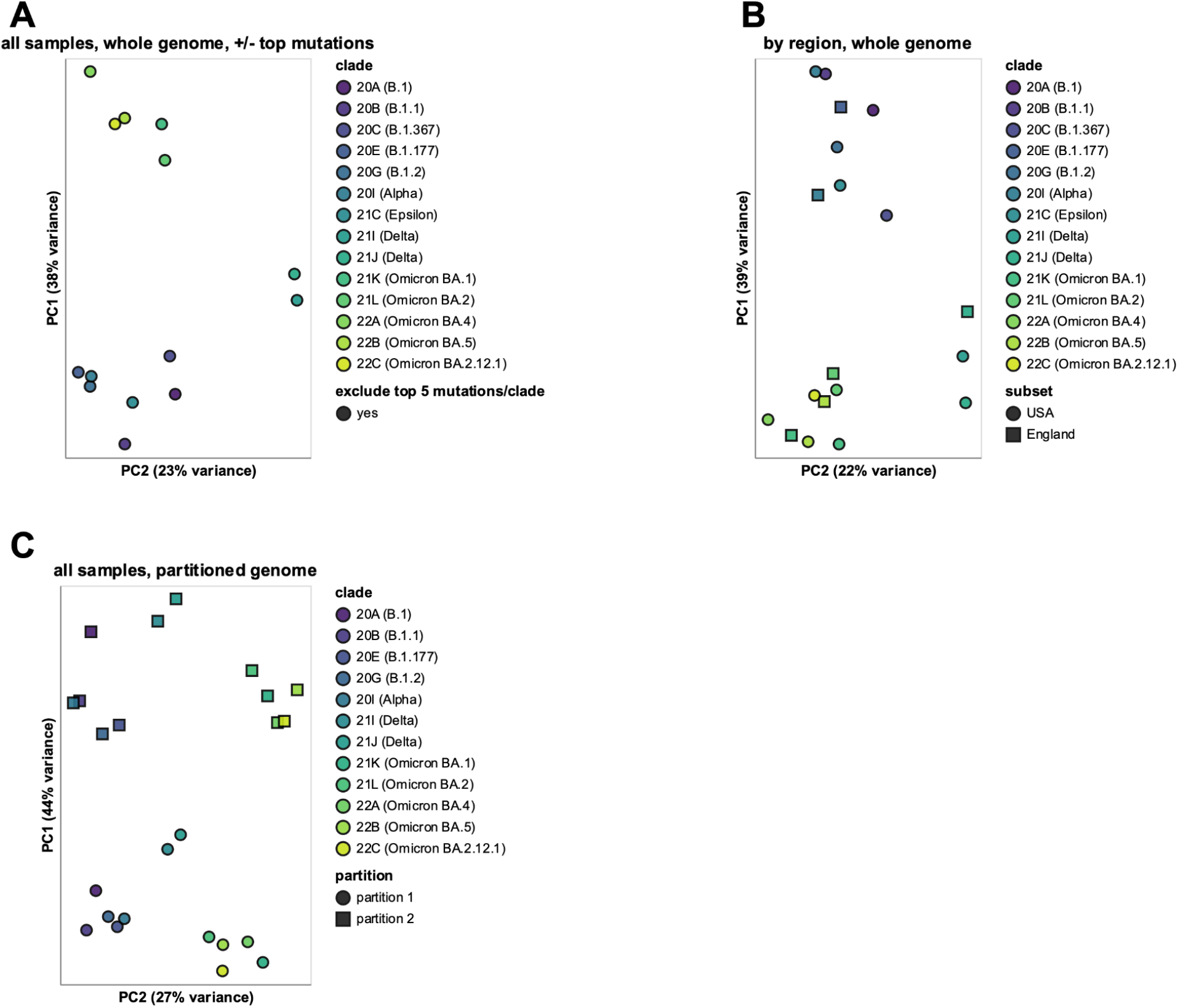
The inter-clade differences in mutation spectrum are robust to various possible sources of noise. This figure repeats the PCA in Figure S1 and shows that the results are robust to (A) excluding sites of the top-5 most abundant mutations in each clade, (B) examining only sequences from the USA or England, or (C) partitioning the genome into half. For panel C, some structure in the PCA plot is explainable by the genome partitioning but the shift in points caused by partitioning the genome is consistent across all clades, and so is not responsible for the inter-clade differences. These plots can be more easily explored using the interactive versions at https://jbloomlab.github.io/SARS2-mut-spectrum/ that enable mousing over of points and clicking on the legend to choose specific clades or groupings.

**Figure S2:**
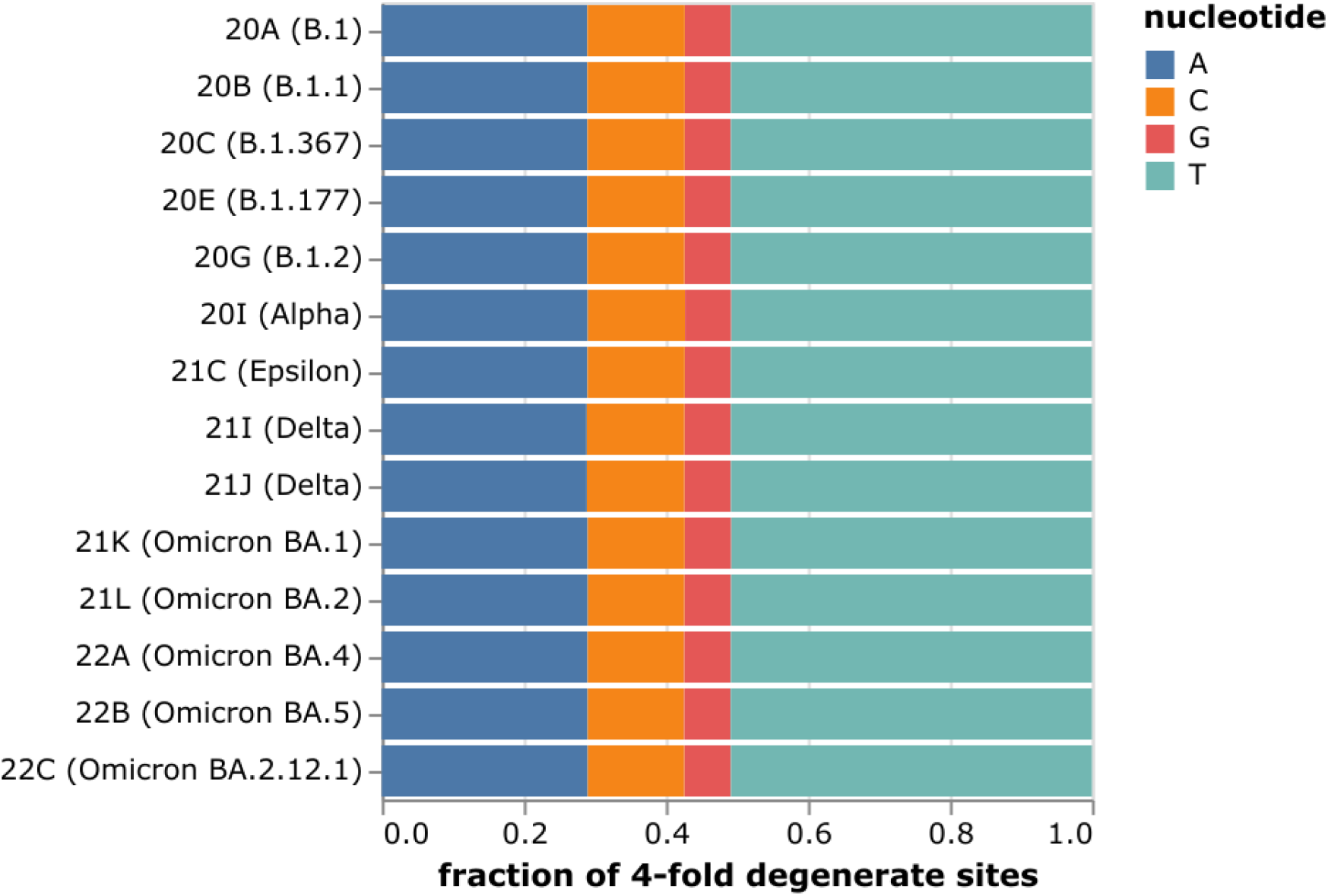
The frequencies of nucleotides at four-fold degenerate sites are nearly identical among the clade founder sequences.

## References

Aksamentov I, Roemer C, Hodcroft EB, Neher RA. 2021. Nextclade: clade assignment, mutation calling and quality control for viral genomes. J. Open Source Softw. 6:3773.

Bessa LM, Guseva S, Camacho-Zarco AR, Salvi N, Maurin D, Perez LM, Botova M, Malki A, Nanao M, Jensen MR, et al. 2022. The intrinsically disordered SARS-CoV-2 nucleoprotein in dynamic complex with its viral partner nsp3a. Sci. Adv. 8:eabm4034.

Cao Y, Jian F, Wang J, Yu Y, Song W, Yisimayi A, Wang J, An R, Chen X, Zhang N, et al. 2022. Imprinted SARS-CoV-2 humoral immunity induces convergent Omicron RBD evolution. :2022.09.15.507787. Available from: https://www.biorxiv.org/content/10.1101/2022.09.15.507787v4

Cao Z, Xia H, Rajsbaum R, Xia X, Wang H, Shi P-Y. 2021. Ubiquitination of SARS-CoV-2 ORF7a promotes antagonism of interferon response. Cell. Mol. Immunol. 18:746–748.

Couce A, Guelfo JR, Blázquez J. 2013. Mutational Spectrum Drives the Rise of Mutator Bacteria. PLOS Genet. 9:e1003167.

De Maio N, Walker CR, Turakhia Y, Lanfear R, Corbett-Detig R, Goldman N. 2021. Mutation Rates and Selection on Synonymous Mutations in SARS-CoV-2. Genome Biol. Evol. 13:evab087.

Denison MR, Graham RL, Donaldson EF, Eckerle LD, Baric RS. 2011. Coronaviruses: an RNA proofreading machine regulates replication fidelity and diversity. RNA Biol. 8:270–279.

Drake JW. 1993. Rates of spontaneous mutation among RNA viruses. Proc. Natl. Acad. Sci. 90:4171–4175.

Eckerle LD, Becker MM, Halpin RA, Li K, Venter E, Lu X, Scherbakova S, Graham RL, Baric RS, Stockwell TB, et al. 2010. Infidelity of SARS-CoV Nsp14-exonuclease mutant virus replication is revealed by complete genome sequencing. PLoS Pathog. 6:e1000896.

Eckerle LD, Lu X, Sperry SM, Choi L, Denison MR. 2007. High Fidelity of Murine Hepatitis Virus Replication Is Decreased in nsp14 Exoribonuclease Mutants. J. Virol. 81:12135–12144.

Felsenstein J. 2003. Inferring Phylogenies. Available from: https://www.amazon.com/Inferring-Phylogenies-Joseph-Felsenstein/dp/0878931775

Fung S-Y, Siu K-L, Lin H, Chan C-P, Yeung ML, Jin D-Y. 2022. SARS-CoV-2 NSP13 helicase suppresses interferon signaling by perturbing JAK1 phosphorylation of STAT1. Cell Biosci. 12:36.

Goldberg and Harris. 2022. Mutational signatures of replication timing and epigenetic modification persist through the global divergence of mutation spectra across the great ape phylogeny. Genome Biol. Evol. 14.

Hardy OJ, Pavoine S. 2012. Assessing phylogenetic signal with measurement error: a comparison of Mantel tests, Blomberg et al.’s K, and phylogenetic distograms. Evol. Int. J. Org. Evol. 66:2614–2621.

Harmon LJ, Glor RE. 2010. Poor statistical performance of the Mantel test in phylogenetic comparative analyses. Evol. Int. J. Org. Evol. 64:2173–2178.

Harris K. 2015. Evidence for recent, population-specific evolution of the human mutation rate. Proc. Natl. Acad. Sci. 112:3439–3444.

Hisner R. 2022. Sublineage of BM.2 with 8 additional spike mutations (9 seq, Australia) · Issue #1286 · cov-lineages/pango-designation. GitHub [Internet]. Available from: https://github.com/cov-lineages/pango-designation/issues/1286

Huerta-Cepas J, Serra F, Bork P. 2016. ETE 3: Reconstruction, Analysis, and Visualization of Phylogenomic Data. Mol. Biol. Evol. 33:1635–1638.

Hwang DG, Green P. 2004. Bayesian Markov chain Monte Carlo sequence analysis reveals varying neutral substitution patterns in mammalian evolution. Proc. Natl. Acad. Sci. 101:13994–14001.

Jiang P, Ollodart AR, Sudhesh V, Herr AJ, Dunham MJ, Harris K. 2021. A modified fluctuation assay reveals a natural mutator phenotype that drives mutation spectrum variation within Saccharomyces cerevisiae. Nordborg M, Przeworski M, editors. eLife 10:e68285.

Kaplanis J, Ide B, Sanghvi R, Neville M, Danecek P, Coorens T, Prigmore E, Short P, Gallone G, McRae J, et al. 2022. Genetic and chemotherapeutic influences on germline hypermutation. Nature. 605:503–508.

Kirchdoerfer RN, Ward AB. 2019. Structure of the SARS-CoV nsp12 polymerase bound to nsp7 and nsp8 co-factors. Nat. Commun. 10:2342.

Legendre P, Legendre L. 2012. Numerical Ecology, Volume 24 - 3rd Edition. Available from: https://www.elsevier.com/books/numerical-ecology/legendre/978-0-444-53868-0

Lindsay, et al. 2019. Similarities and differences in patterns of germline mutation between mice and humans. Nat. Commun. 10.

Liu Y, Qin C, Rao Y, Ngo C, Feng JJ, Zhao J, Zhang S, Wang T-Y, Carriere J, Savas AC, et al. 2021. SARS-CoV-2 Nsp5 Demonstrates Two Distinct Mechanisms Targeting RIG-I and MAVS To Evade the Innate Immune Response. mBio 12:e02335–21.

Long H, Kucukyildirim S, Sung W, Williams E, Lee H, Ackerman M, Doak TG, Tang H, Lynch M. 2015. Background Mutational Features of the Radiation-Resistant Bacterium Deinococcus radiodurans. Mol. Biol. Evol. 32:2383–2392.

Macià MC, Skov L, Peter BM, Schierup MH. 2021. Different historical generation intervals in human populations inferred from Neanderthal fragment lengths and mutation signatures. Nat. Commun. [Internet] 12. Available from: https://www.ncbi.nlm.nih.gov/pmc/articles/PMC8423828/

Mantel. 1967. Ranking Procedures for Arbitrarily Restricted Observation. Biometrics 23:65–78.

Martin DP, Weaver S, Tegally H, San JE, Shank SD, Wilkinson E, Lucaci AG, Giandhari J, Naidoo S, Pillay Y, et al. 2021. The emergence and ongoing convergent evolution of the SARS-CoV-2 N501Y lineages. Cell 184:5189–5200.e7.

Mathieson I, Reich D. 2017. Differences in the rare variant spectrum among human populations. PLOS Genet. 13:e1006581.

Minh BQ, Schmidt HA, Chernomor O, Schrempf D, Woodhams MD, Haeseler A von, Lanfear R. 2020. IQ-TREE 2: New Models and Efficient Methods for Phylogenetic Inference in the Genomic Era. Mol. Biol. Evol. 37:1530.

Mölder F, Jablonski KP, Letcher B, Hall MB, Tomkins-Tinch CH, Sochat V, Forster J, Lee S, Twardziok SO, Kanitz A, et al. 2021. Sustainable data analysis with Snakemake. F1000Research 10:33.

Neher RA. 2022. Contributions of adaptation and purifying selection to SARS-CoV-2 evolution. :2022.08.22.504731. Available from: https://www.biorxiv.org/content/10.1101/2022.08.22.504731v1

Ogando NS, Zevenhoven-Dobbe JC, van der Meer Y, Bredenbeek PJ, Posthuma CC, Snijder EJ. 2020. The Enzymatic Activity of the nsp14 Exoribonuclease Is Critical for Replication of MERS-CoV and SARS-CoV-2. J. Virol. 94:e01246–20.

Oksanen J, Simpson GL, Blanchet FG, Kindt R, Legendre P, Minchin PR, O’Hara RB, Solymos P, Stevens MHH, Szoecs E, et al. 2022. vegan: Community Ecology Package. Available from: https://CRAN.R-project.org/package=vegan

Pauly MD, Lyons DM, Fitzsimmons WJ, Lauring AS. 2017. Epistatic Interactions within the Influenza A Virus Polymerase Complex Mediate Mutagen Resistance and Replication Fidelity. mSphere 2:e00323–17.

Peck KM, Lauring AS. 2018. Complexities of Viral Mutation Rates. J. Virol. 92:e01031–17.

Pickering B, Lung O, Maguire F, Kruczkiewicz P, Kotwa JD, Buchanan T, Gagnier M, Guthrie JL, Jardine CM, Marchand-Austin A, et al. 2022. Highly divergent white-tailed deer SARS-CoV-2 with potential deer-to-human transmission. :2022.02.22.481551. Available from: https://www.biorxiv.org/content/10.1101/2022.02.22.481551v1

Ratcliff J, Simmonds P. 2021. Potential APOBEC-mediated RNA editing of the genomes of SARS-CoV-2 and other coronaviruses and its impact on their longer term evolution. Virology 556:62–72.

Ringlander J, Fingal J, Kann H, Prakash K, Rydell G, Andersson M, Martner A, Lindh M, Horal P, Hellstrand K, et al. 2022. Impact of ADAR-induced editing of minor viral RNA populations on replication and transmission of SARS-CoV-2. Proc. Natl. Acad. Sci. 119:e2112663119.

Robinson, et al. 2021. Increased somatic mutation burdens in normal human cells due to defective DNA polymerases. Nat. Genet. 53:1434–1442.

Roe MK, Junod NA, Young AR, Beachboard DC, Stobart CC. 2021. Targeting novel structural and functional features of coronavirus protease nsp5 (3CLpro, Mpro) in the age of COVID-19. J. Gen. Virol. 102.

Ruis C, Peacock TP, Polo LM, Masone D, Alvarez MS, Hinrichs AS, Turakhia Y, Cheng Y, McBroome J, Corbett-Detig R, et al. 2022. Mutational spectra distinguish SARS-CoV-2 replication niches. :2022.09.27.509649. Available from: https://www.biorxiv.org/content/10.1101/2022.09.27.509649v1

Ruis C, Weimann A, Tonkin-Hill G, Pandurangan AP, Matuszewska M, Murray GGR, Lévesque RC, Blundell TL, Floto RA, Parkhill J. 2022. Mutational spectra analysis reveals bacterial niche and transmission routes. :2022.07.13.499881. Available from: https://www.biorxiv.org/content/10.1101/2022.07.13.499881v1

Sadler HA, Stenglein MD, Harris RS, Mansky LM. 2010. APOBEC3G Contributes to HIV-1 Variation through Sublethal Mutagenesis. J. Virol. 84:7396–7404.

Sasani TA, Ashbrook DG, Beichman AC, Lu L, Palmer AA, Williams RW, Pritchard JK, Harris K. 2022. A natural mutator allele shapes mutation spectrum variation in mice. Nature 605:497–502.

Speidel L, Cassidy L, Davies RW, Hellenthal G, Skoglund P, Myers SR. 2021. Inferring Population Histories for Ancient Genomes Using Genome-Wide Genealogies. Mol. Biol. Evol. 38:3497–3511.

Sung W, Ackerman MS, Miller SF, Doak TG, Lynch M. 2012. Drift-barrier hypothesis and mutation-rate evolution. Proc. Natl. Acad. Sci. U. S. A. 109:18488–18492.

Sung W, Tucker AE, Doak TG, Choi E, Thomas WK, Lynch M. 2012. Extraordinary genome stability in the ciliate Paramecium tetraurelia. Proc. Natl. Acad. Sci. 109:19339–19344.

Turakhia Y, Maio ND, Thornlow B, Gozashti L, Lanfear R, Walker CR, Hinrichs AS, Fernandes JD, Borges R, Slodkowicz G, et al. 2020. Stability of SARS-CoV-2 phylogenies. PLOS Genet. 16:e1009175.

Turakhia Y, Thornlow B, Hinrichs AS, De Maio N, Gozashti L, Lanfear R, Haussler D, Corbett-Detig R. 2021. Ultrafast Sample placement on Existing tRees (UShER) enables real-time phylogenetics for the SARS-CoV-2 pandemic. Nat. Genet. 53:809–816.

VanderPlas J, Granger BE, Heer J, Moritz D, Wongsuphasawat K, Satyanarayan A, Lees E, Timofeev I, Welsh B, Sievert S. 2018. Altair: Interactive Statistical Visualizations for Python. J. Open Source Softw. 3:1057.

Vignuzzi M, Wendt E, Andino R. 2008. Engineering attenuated virus vaccines by controlling replication fidelity. Nat. Med. 14:154–161.

V’kovski P, Kratzel A, Steiner S, Stalder H, Thiel V. 2021. Coronavirus biology and replication: implications for SARS-CoV-2. Nat. Rev. Microbiol. 19:155–170.

Zhang J, Ejikemeuwa A, Gerzanich V, Nasr M, Tang Q, Simard JM, Zhao RY. 2022. Understanding the Role of SARS-CoV-2 ORF3a in Viral Pathogenesis and COVID-19. Front. Microbiol. [Internet] 13. Available from: https://www.ncbi.nlm.nih.gov/pmc/articles/PMC8959714/

Ziebuhr J. 2005. The Coronavirus Replicase. Coronavirus Replication Reverse Genet. 287:57–94.

